# DPSN: standardizing the short names of amplicon-sequencing primers to avoid ambiguity

**DOI:** 10.1101/2020.01.23.916429

**Authors:** Yuxiang Tan, Yixia Tian, Junyu Chen, Zhinan Yin, Hengwen Yang

**Affiliations:** The First Affiliated Hospital, Biomedical Translational Research Institute, Guangdong Province Key Laboratory of Molecular Immunology and Antibody Engineering, Jinan University, Guangzhou 510632, China; Shenzhen Institute of Synthetic Biology, Shenzhen Institutes of Advanced Technology, Chinese Academy of Sciences, Shenzhen 518055, People’s Republic of China

**Keywords:** database, scientific name, amplicon, primer

## Abstract

**Background:** Amplicon sequencing is the most widely used sequencing method to evaluate microbial diversity in virtually all environments. Thus, appropriate and specific primers are needed to amplify amplicon regions in amplicon sequencing. For this purpose, the community currently uses probeBase, which curates rRNA-targeted probes and primers.

**Main Body:** We found that 63.58% of the primers in probeBase have problematic issues in the short name, full name, and/or position. Furthermore, the current convention for short names causes ambiguity. We here introduce our new Database of Primer Scientific Names (DPSN), which is a manually curated database for the 173 primers in probeBase complete with a new short name convention. Building on the work of probeBase, we provide a more user-friendly and standardized system. The new short primer naming convention has three basic components: 5□ position on the sense strand, version, and direction. An additional character for the name of the taxonomic group is also added in front of the name for convenient use. Furthermore, DPSN contains primers for large subunit as well. In order to separate them from the primers for small subunit, a header character is also recommended.

**Conclusion:** All 173 primers in probeBase were corrected according to this new rule, and are stored in DPSN, which is expected to facilitate accurate primer selection and better standardized communication in this field.

**Database URL:** The DPSN database is available in a user-interactive website at http://dpsn.leidailab.cn/

## Background

Amplicon sequencing is a common sequencing method for microbial research from diverse environmental or clinical samples [1, 2]. Amplicon sequencing is dependent on the choice of primers for carrying out the amplification step. Thus, selection of the most appropriate primers is the foundation of successful amplicon sequencing.

probeBase [3] is the only database currently available with updated lists of probes and primers, along with links to other databases providing related information. At present, there is a total of 173 primers recorded in probeBase. In general, a primer is defined according to its short name (SN), full name (FN), and sequence. However, in many cases, only the SN is used for the sake of convenience. There are seven components of an FN [4]. Taking S-D-Bact-0338-a-A-18 as an example: “S” stands for the target gene (Small Sub-Unit (SSU)), “D” represents the largest taxonomic group targeted (Domain), “Bact” is the name of the taxonomic group (Bacteria), “0338” is the 5□ position of the sense strand, “a” presents the version, “A” denotes the identical strand (“S” for sense; “A” for antisense), and “18” is the length of the primer. To avoid ambiguity, each primer should have a unique SN; however, this is not the case. Different from FN, there is no guideline for how an SN should be. Therefore, SNs were named in a few different ways, such as Primer3, Bac927, 926r, and 934mcr. The most common ones were composited by the position and direction (for example, 926r), or with an additional string for the name of the taxonomic group (for example, Arch 915r). The lack of clear rules and sufficient information for accuracy leads to ambiguity of SNs.

In fact, there are 14 SNs that refer to multiple primer sequences, which could lead to confusion and cause several problems in application for users. For example, in the earth microbiome project website [1], the author of the citation for a given primer is used along with the SN to better specify the primer. Furthermore, the SN itself could be misleading. For example, primers 907r and 926r are actually from the same region of the genome but with a difference of two bases in the sequence. However, based on their SNs alone, a user would misinterpret these primers as being derived from two different regions.

To resolve this problem, we here introduce Database of Primer Scientific Names (DPSN), which is a database that has been manually curated to correct problematic and inconsistent features (SN, FN, position, and length) of primers in probeBase according to an improved convention of naming SNs. The new SNs still correspond to the old SNs and corrected FNs in a one-to-one relation.

## Construction and content

### Data source

Information of all 173 primers in the probeBase dataset was manually extracted [3], including the SN, FN, position, sequence, length, G+C content, and dissociation temperature.

The corresponding regions on the reference sequence of *Escherichia coli* K-12 substrain MG1655 was extracted from the SILVA database [5] and served as the reference for confirming the sequence position.

### Derivation of new SN naming convention

A unique SN should have at least three basic components to provide sufficient identifying information: 5□ position on the sense strand, version, and direction.

In the old SN, such as 907r, all reverse primers that start or end from position 907 will have the same name, which leads to ambiguity. Therefore, including additional information of the version, such as 907ar to indicate the version, could help to specify the primer sequence. Consistent with the old SN rule, “f” and “r” denote “forward primer” and “reverse primer,” respectively. Moreover, because the name of the taxonomic group provides helpful information for users to select appropriate primers, DPSN also includes a shorthand for the name of the taxonomic group (“A” for Archaea, “B” for Bacteria, “U” for universal, and “N” for nano) in front of the SN. Additionally, on account of the need to separate SSU primers from large subunit (LSU) primers, the header represents of target get from the FNs is retained. For instance, the SN 907ar represents the FN “S-D-Bact-0907-a-A-20”, which was corrected and recorded as S-B907ar in DPSN.

### Amplification range validation of the primers

To validate the amplification location of primers according to the *E. coli* K-12 reference, BLAT of the National Center for Biotechnology Information [6] was employed as the aligner. However, because of the presence of degenerate bases in primer sequences, the primers had to be converted into expanded regular sequences, which was achieved using a customized Python script before BLAT alignment. In particular, the additional parameters “-minMatch = 1-minScore = 8-minIdentity = 70-stepSize =1-tileSize =8” were applied to BLAT, considering the length and mismatch tolerance of primers.

To confirm the amplification location, the primers were also checked by the TestProbe function in SILVA [5].

## Utility and Discussion

### How to use DPSN

In order to make the search easy, DPSN supports fuzzy search on all the fields. This means user can use any keyword to find the related primer(s), such as the intended 5’ position and the sub-string of the primer sequence. In return, DPSN will present the corrected information of the related primers. As well as the original “Short Name” and “Full Name” from the probeBase, which will help the user to connect the use of the primers in original papers. All the sequence in DPSN are the same as the ones in probeBase.

### Summary of Corrections and Discussion

Our careful review of probeBase identified five sequences with multiple primer names, 14 groups of SNs with multiple targets, and 91 SNs inconsistent with their FNs. Of the total 173 primers in probeBase, the SNs for only 63 primers (36.42%) could direct the user to a unique sequence and be considered as correct. Five sequences were multifold and had multiple primer names (Table 1). Thirty SNs from 14 groups pointed to more than one sequence. The positions in 91 SNs were different from the 5□ position of their FNs. Overall, the positions of 35 primers in probeBase were found to be incorrect.

**Table 1:**
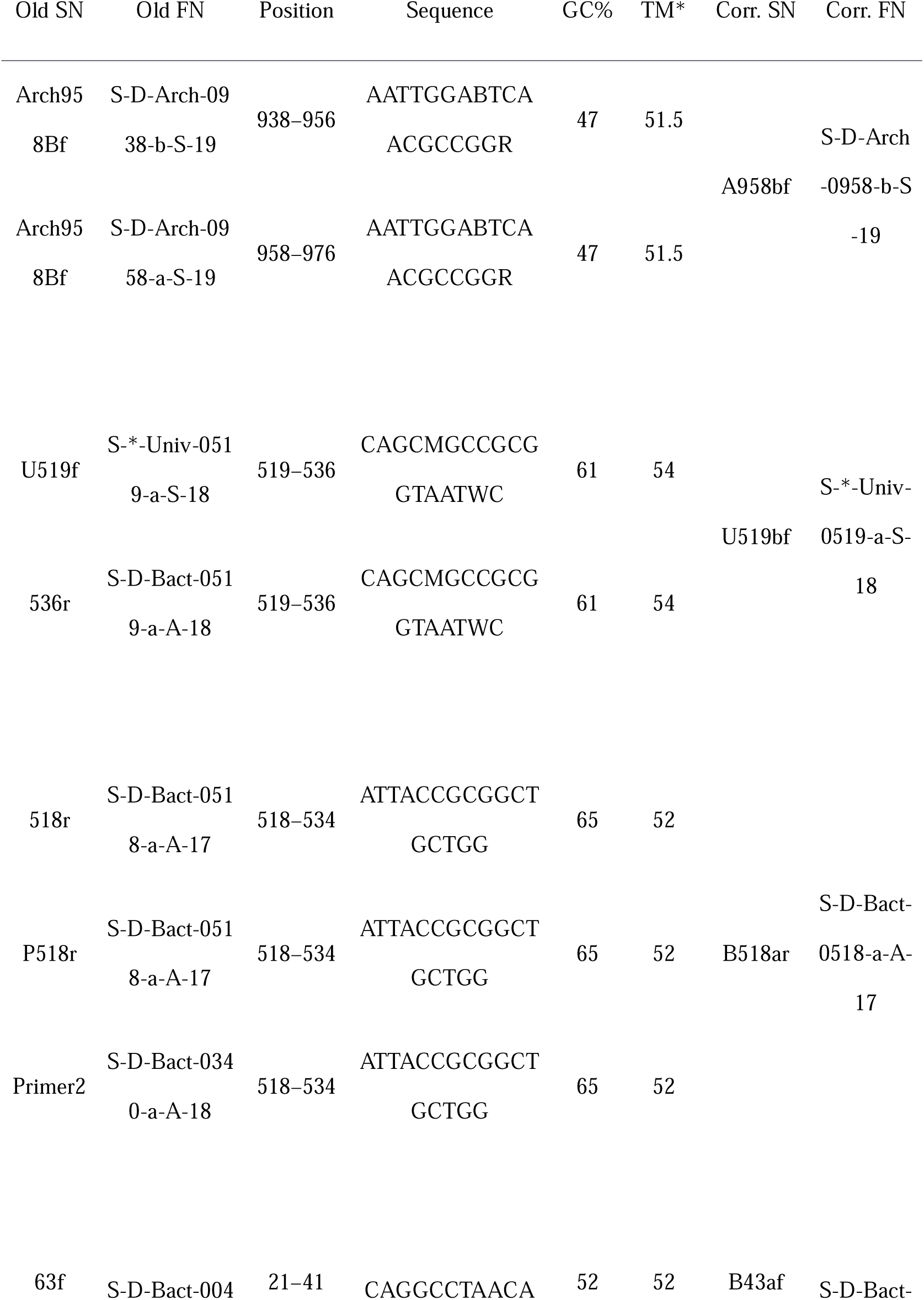
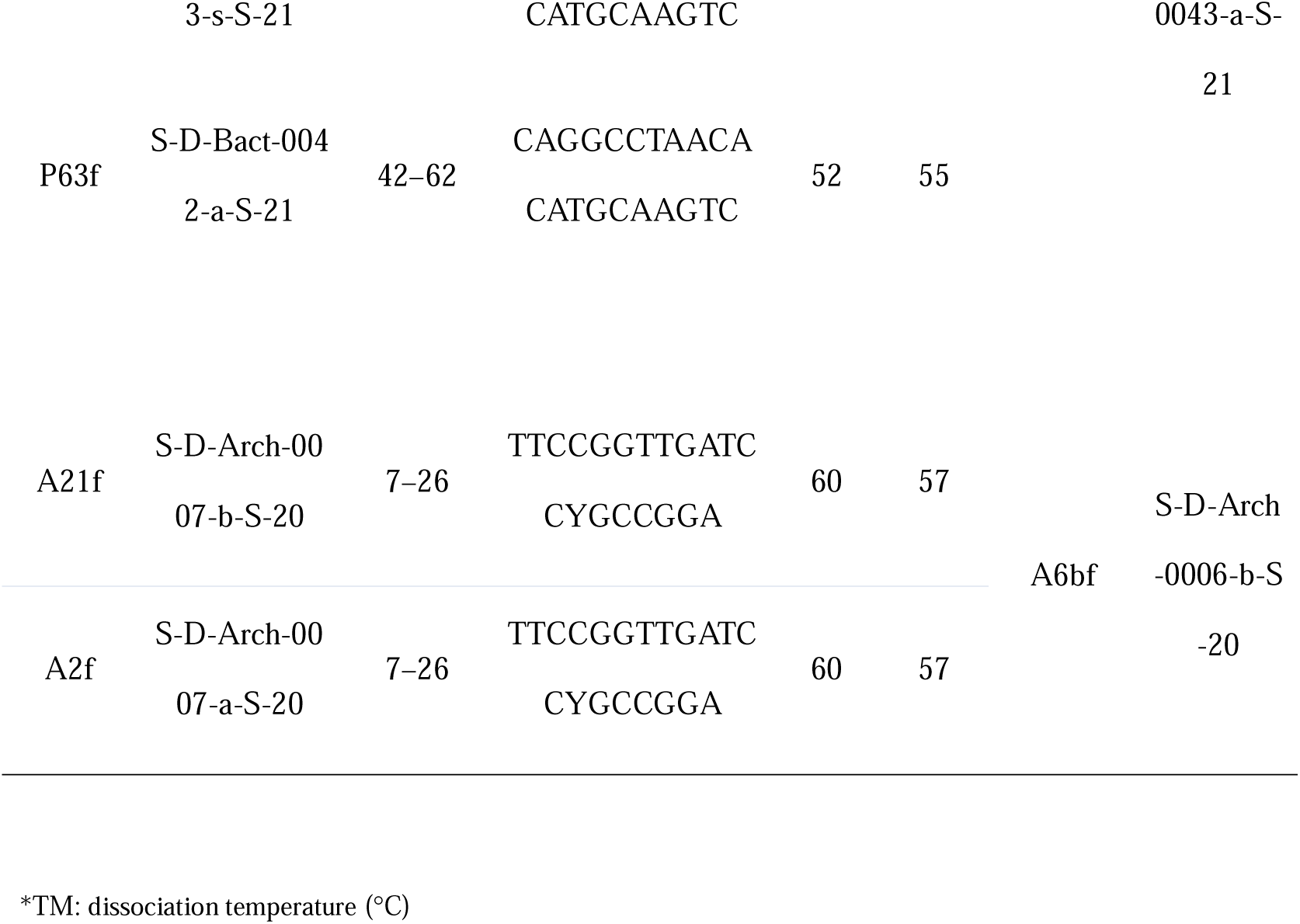
Primer groups with the same sequence in probeBase.

In addition, a few FNs were found to be incorrect in probeBase, which have been manually corrected in DPSN. Theoretically, the FNs of primers should be unique, since a single FN represents a unique primer sequence. However, in probeBase, three FNs were duplicated and even represented more than one sequence (Table 2). In the naming rule, the position in an FN is based on the 5□ position; however, eight of the primers in probeBase violated this rule (Table 3). Even more importantly, the length information of five FNs did not match the actual lengths of their sequences (Table 4), and the directions of three primers were opposite to the actual direction of their sequences (Table 5).

**Table 2:**
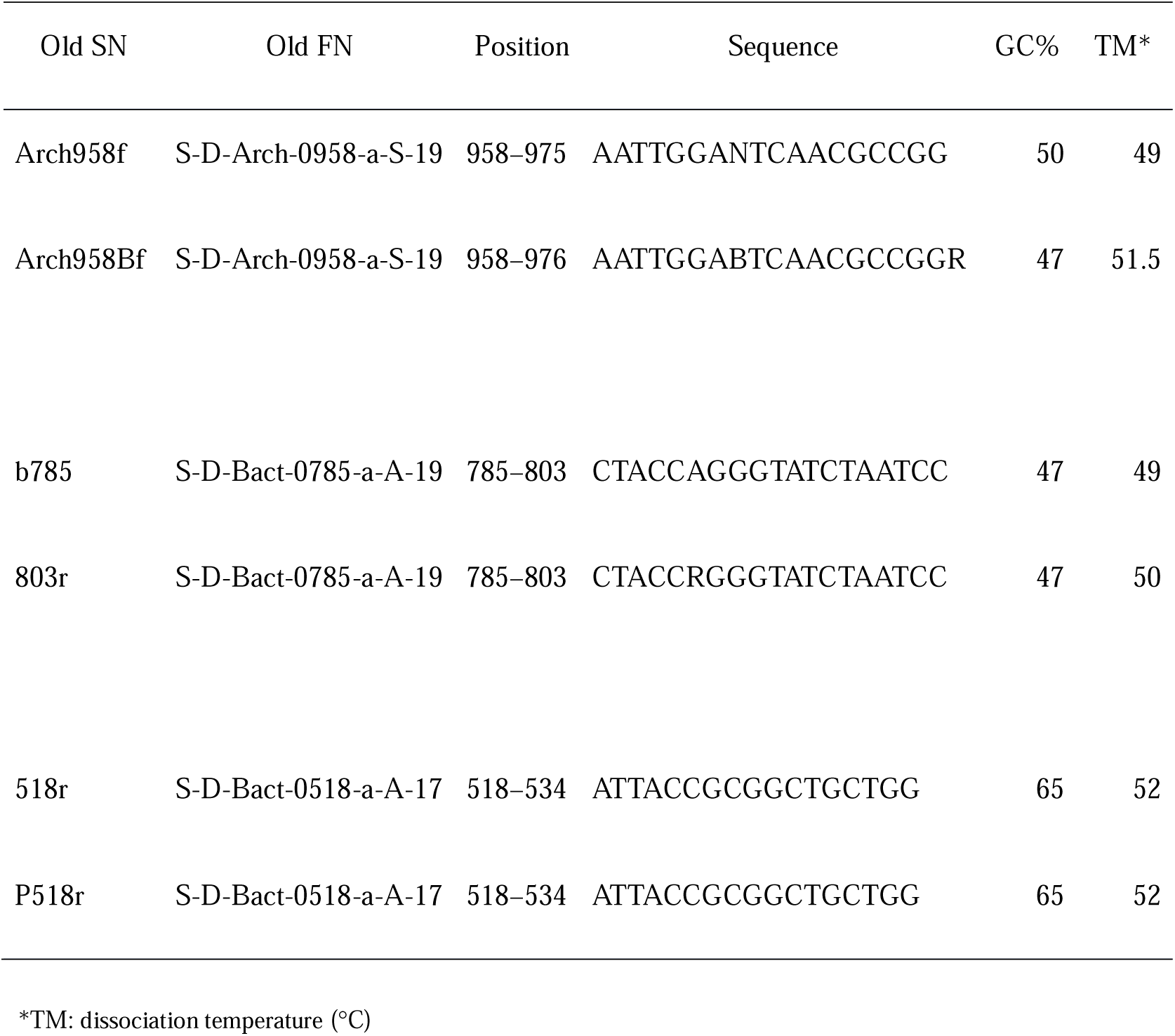
Primer groups with the same FN in probeBase.

**Table 3:**
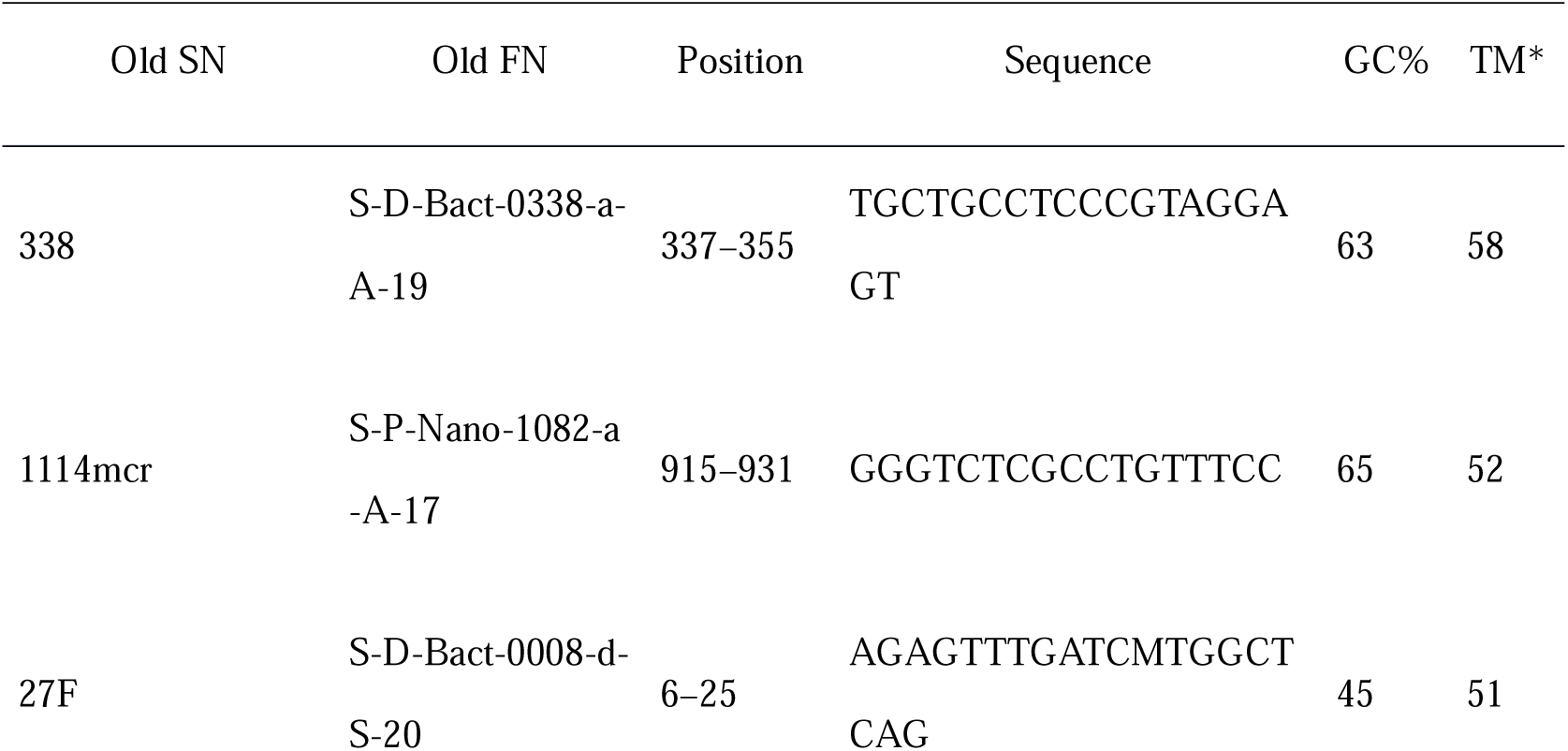
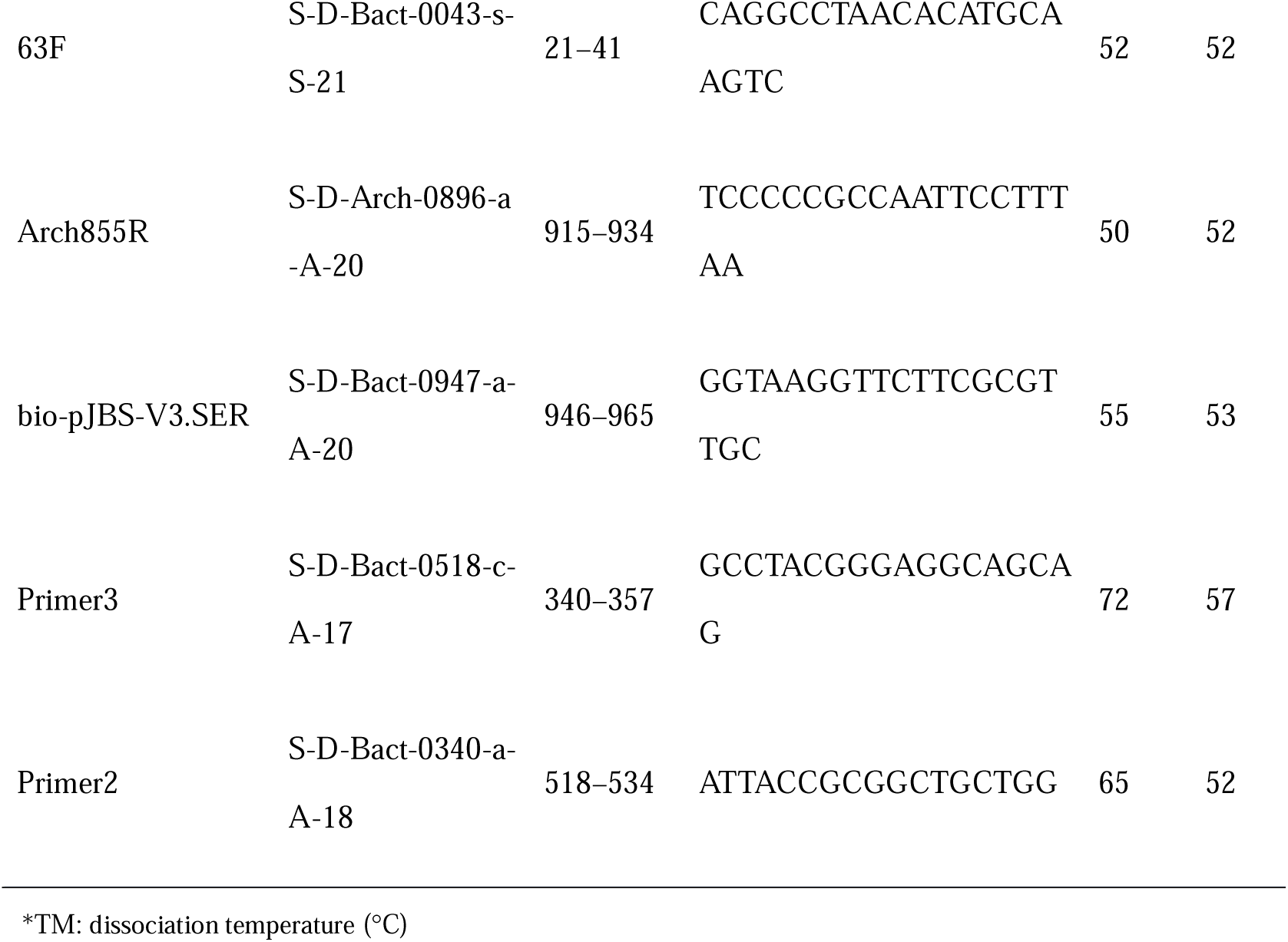
FNs of primers with inconsistent positions in probeBase.

**Table 4:**
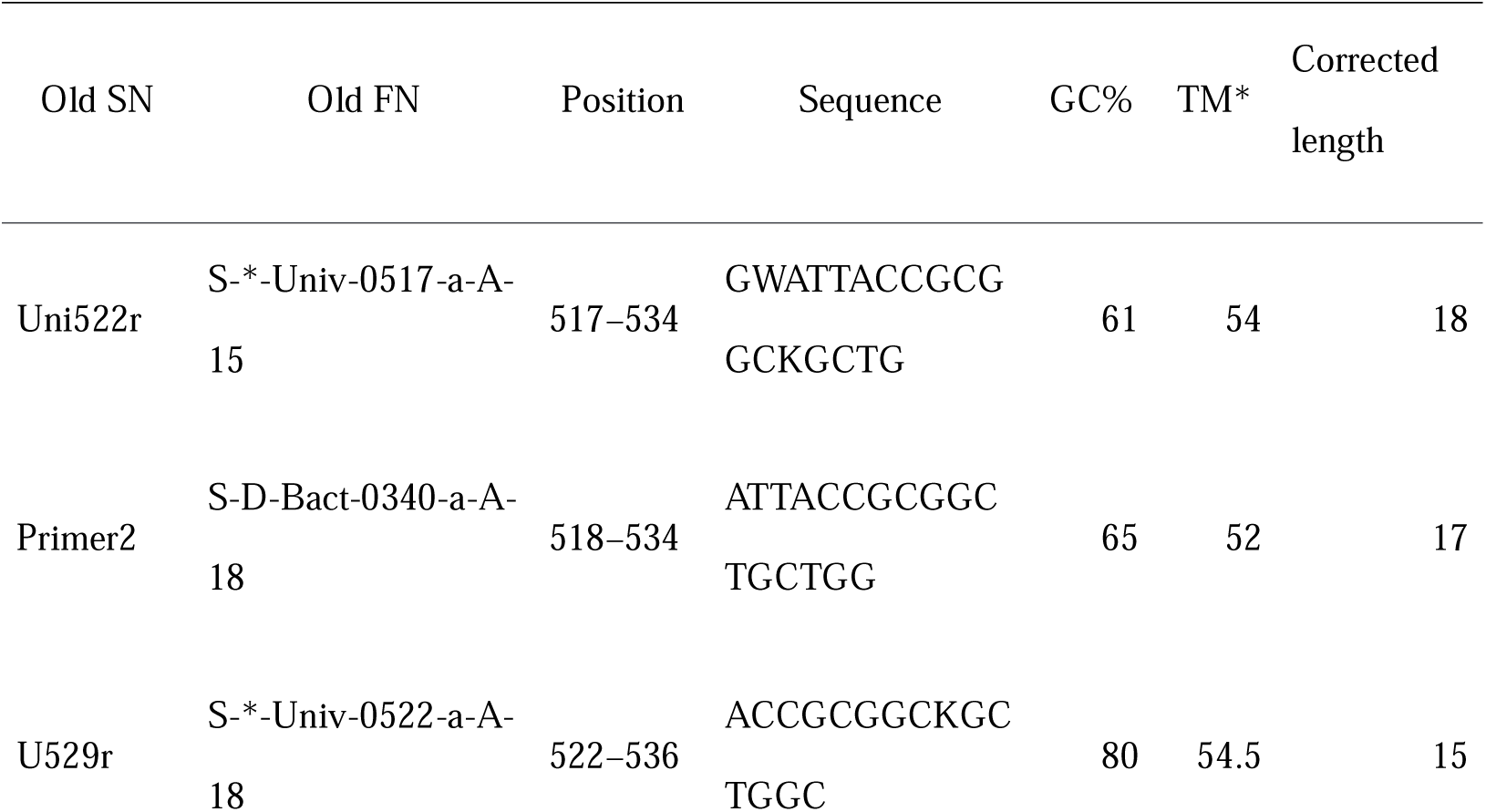
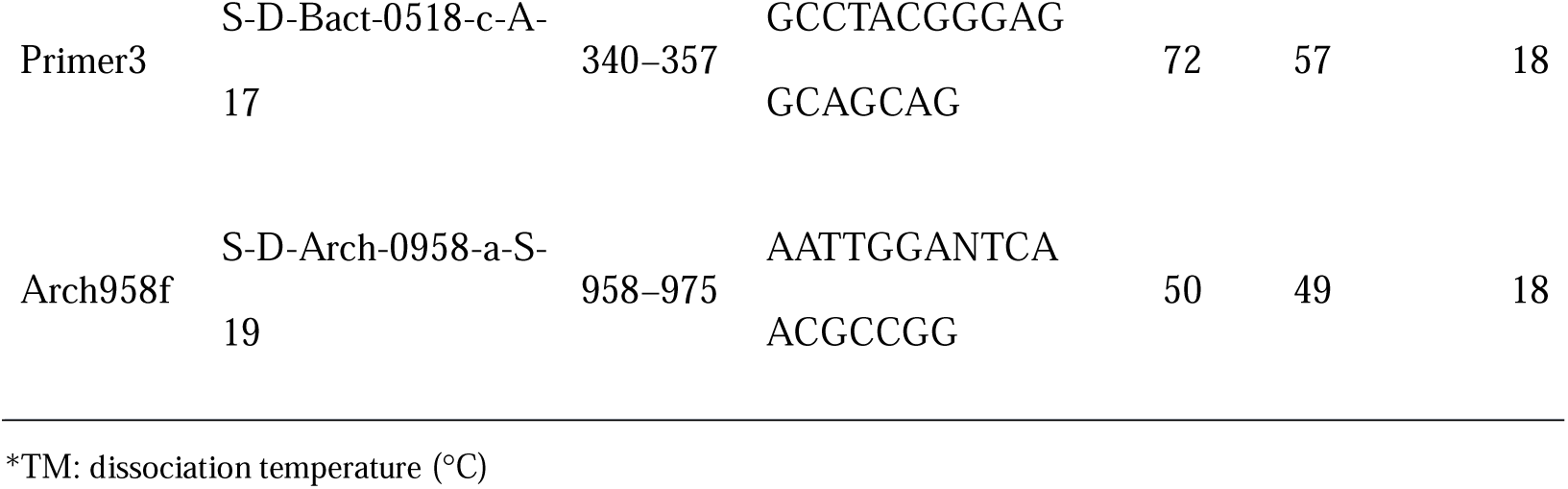
FNs of primers with the wrong length in probeBase.

**Table 5:**
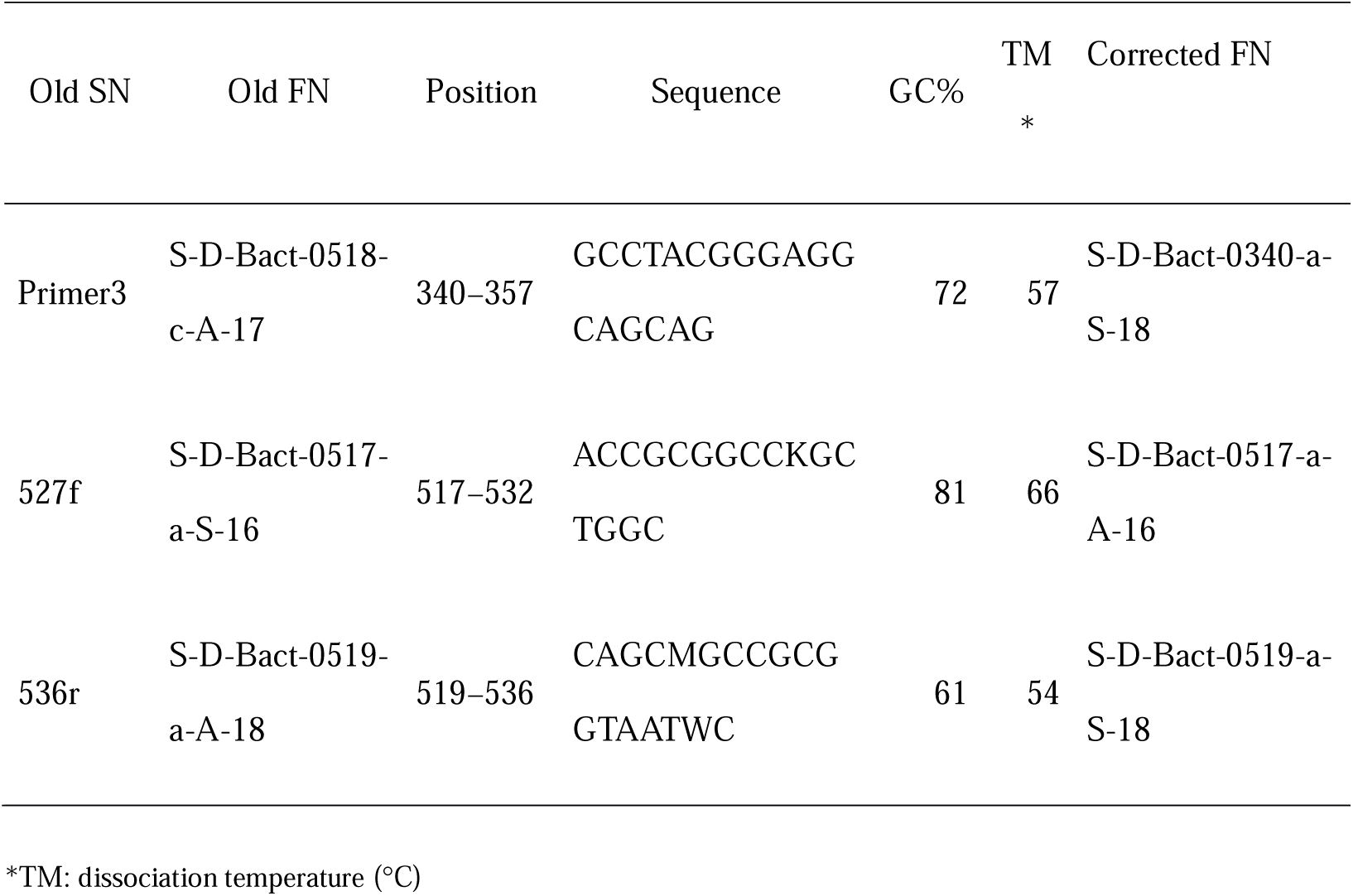
FNs of primers with the wrong strand in probeBase.

In DPSN, all of the SNs of the primers in probeBase have been updated according to the new naming rule along with additional version information to provide a more unique identifier that is still convenient to use. Overall, 110 problematic primers were corrected. Using the amended primer name in DPSN, users can simply refer to the SN to specify a primer, because of the one-to-one relation among the SN, FN, and sequence, and without the inconvenience of appending additional information such as author name or sequence in the article.

## Conclusion

In conclusion, because it is crucial to avoid vagueness in scientific research, the old SN system of primers is problematic and should be replaced by the new naming rule as proposed herein. All of these corrections have been curated in DPSN to improve searching convenience and accuracy. Therefore, with DPSN, users can easily search an old name from probeBase or articles for its amended name. For new articles, it is recommended that authors use the amended name to accurately describe a primer.

DPSN currently focuses on only primers for amplicon sequencing on SSU and LSU, and thus it can be assumed that the ambiguity problem still exists for primers in other amplicon regions, such as ITS. Because the primers for these regions were not found in probeBase, we can collect and import these primers into the naming system in DPSN in the future. To keep the database up to date, DPSN accepts data submission of primers from researchers as well.

## Abbreviations

DPSN: Database of Primers’ Scientific Names
SN: short name
FN: full name

## Declarations

### Ethics approval and consent to participate

Not applicable

### Consent for publication

Not applicable

### Availability of data and material

<u>Database Name:</u> Database of Primers’ Scientific Names (DPSN)

<u>Database URL:</u> http://dpsn.leidailab.cn/

### Authors’ contributions

YuT conceived of the idea, conducted the analysis, and wrote the manuscript. YiT performed the data collection. JC provided data of LSU primers. HY and ZY supervised the project and participated in the revision of the manuscript. All authors read and approved the final manuscript.

## Acknowledgements

We would like to thank Becky Kusko for editing suggestions and Editage (www.editage.com) for English language editing.

## References

1. Gilbert JA, Jansson JK, Knight R. The Earth Microbiome project: successes and aspirations. BMC Biol. 2014;12:69.

2. The Human Microbiome Project Consortium. A framework for human microbiome research. Nature 2012;486:215–21.

3. Greuter D, Loy A, Horn M, Rattei T. probeBase—an online resource for rRNA-targeted oligonucleotide probes and primers: new features. Nucleic Acids Res. 2016;44:D586–9.

4. Klindworth A, Pruesse E, Schwwer T, Peplies J, Quast C, Horn M, et al. Evaluation of general 16S ribosomal RNA gene PCR primers for classical and next-generation sequencing-based diversity studies. Nucleic Acids Res. 2013;41:e1.

5. Quast C, Pruesse E, Yilmaz P, Gerken J, Schwwer T, Yarza P, Peplies J, Glöckner FO. The SILVA ribosomal RNA gene database project: improved data processing and web-based tools. Nucleic Acids Res. 2013;41:D590–6.

6. Kent WJ. BLAT--the BLAST-like alignment tool. Genome Res. 2002;12:656–4.

